# Visual semantic tuning across the cortex shifts between tasks

**DOI:** 10.64898/2026.02.19.706797

**Authors:** Tianjiao Zhang, Jack L. Gallant

**Author notes:** Corresponding author: Jack L. Gallant.

## Abstract

Attention is a powerful mechanism that dynamically optimizes brain representations to prioritize behaviorally-relevant information. While previous studies focusing on single tasks have demonstrated that attention shifts tuning towards attended targets, real-world behavior often requires humans to switch between tasks with different demands. How does visual semantic tuning in the human brain shift to support the behavioral demands of different naturalistic tasks? To answer this question, we used voxelwise encoding models to compare visual semantic tuning across the human cerebral cortex between two distinct naturalistic tasks, movie watching and navigation. Results show that visual semantic tuning in the cortex differed substantially between tasks. Principal component analysis reveals that during navigation, tuning shifts increase the representation of vehicles and traffic signs compared to movie watching, and that these shifts were localized to distinct functional networks. These tuning shifts reconfigure the human brain’s representations of object categories based on their behavioral relevance. These findings demonstrate that visual semantic tuning in the brain dynamically shifts towards task-relevant information across naturalistic tasks, optimizing functional representations to achieve diverse behavioral goals.

**Significance statement:** In real-world situations, we switch between many tasks with behaviorally distinct goals that require us to attend to different targets. Here we show that visual semantic tuning across the human cortex shifts between passive movie-watching and active navigation. These tuning shifts increase the representation of behaviorally-relevant objects, such as cars during navigation. Tuning shifts towards different object categories are distributed across distinct functional networks, suggesting that they engage different cognitive mechanisms. These results show that visual semantic tuning in the human brain is highly task-dependent and is optimized to represent behaviorally-relevant information.

## INTRODUCTION

Attention is a powerful mechanism that optimizes brain function to prioritize processing behaviorally-relevant information. Neurophysiology studies in non-human primates have demonstrated that attentional effects are found across the visual hierarchy. Attention modulates the gain of neural firing rates as early as V1 (McAdams and Maunsell, 1999), and also the baseline of neural firing rates in V2 and V4 (Motter, 1993; Luck et al., 1997; McAdams and Maunsell, 1999). Furthermore, spatial attention causes the receptive fields of neurons in both V4 and MT to shift towards the attended location (Connor et al., 1997; Womelsdorf et al., 2006), and feature-based attention shifts the tuning of V4 neurons towards the attended feature (David et al., 2008). Finally, tuning for visual features in prefrontal neurons changes by task context (Asaad et al., 2000; Johnston and Everling, 2006; Warden and Miller, 2010; McKee et al., 2014).

Complementing these neurophysiological findings, neuroimaging studies in humans have found that attention changes voxel tuning across much of the cerebral cortex. Spatial attention shifts population receptive fields across the visual hierarchy (Klein et al., 2014; Vo et al., 2017). Attention to high-level visual features such as object category, action category, or affordances shifts tuning in lateral occipital (Çukur et al., 2013; Kaiser et al., 2016), parietal (Çukur et al., 2013; Bracci et al., 2017), and prefrontal regions (Çukur et al., 2013; Bugatus et al., 2017; Nastase et al., 2017; Shahdloo et al., 2022). Together, these converging neurophysiology and neuroimaging results suggest that attention acts as a matched-filter mechanism that increases the representation of task-relevant information, and which consequently decreases the representation of task-irrelevant information (Mazer and Gallant, 2003; David et al., 2008).

Because different tasks have distinct constraints and behavioral goals, attentional optimization of brain representations is likely to be most prominent when comparing across distinct tasks. Indeed, primate neurophysiology studies have reported that different tasks evoked neuronal tuning shifts in many brain regions, including the inferior temporal cortex (Koida and Komatsu, 2007) and the prefrontal cortex (Asaad et al., 2000; Johnston and Everling, 2006; Warden and Miller, 2010; McKee et al., 2014). However, most human studies have only examined attentional modulation of functional representations within the context of a single task. For example, in a detection task participants might be directed to attend to different screen locations or to search for different object categories, while the general stimulus and task structure remains fixed (Harel et al., 2014; Bracci et al., 2017). Thus, task-related tuning shifts remain poorly studied in humans (Kay et al., 2023).

To better understand tuning shifts across naturalistic tasks in the human brain, we compared visual semantic tuning in the cerebral cortex measured in a movie-watching experiment and a navigation experiment (Fig. 1). Because movie-watching and navigation have very different attentional demands, we hypothesized that there will be significant tuning differences across the cortex between the two tasks. In the movie-watching experiment, participants watched naturalistic videos at fixation (Huth et al., 2012). In the navigation experiment, a different group of participants drove a virtual car from a first-person perspective through a virtual city (Zhang et al., 2025). Brain activity was recorded with fMRI in both experiments. We used voxelwise modelling to characterize the visual semantic tuning across the cortex in both experiments (Naselaris et al., 2011). Comparison of the visual semantic model weights obtained in the movie-watching participants with those obtained from the active navigation task shows that visual semantic tuning differs significantly across the cortex between tasks, and suggest that the tuning shifts serve to enhance the representation of task-relevant categories.

**Figure 1.**
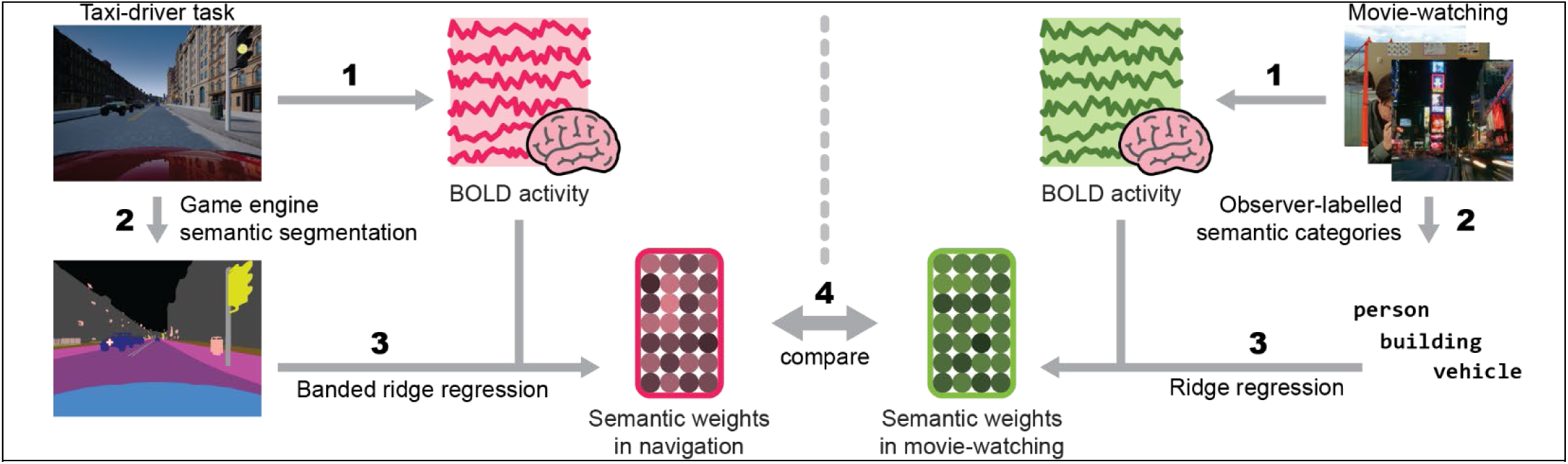
Comparing visual semantic tuning across tasks. Attention is a powerful mechanism for increasing the brain’s efficiency for processing task-relevant information. By operating as a matched-filter mechanism that shifts neural tuning towards targets (Mazer and Gallant, 2003; David et al., 2008), attention dynamically increases the brain’s representation of attentional targets. Because attentional demands differ across task contexts, we hypothesized that visual sematic tuning of the brain will differ between movie-watching and navigation. To test this hypothesis, we compared visual semantic tunings of the brain between a navigation experiment and a movie-watching experiment. In the movie-watching experiment, participants watched clips of various naturalistic movies and videos. To create visual semantic labels for the movies, each scene was semantically labeled by two independent observers. In the navigation experiment, participants drove through a large virtual city from a first-person perspective, and were cued to drive to different locations in the virtual city. To create visual semantic labels for the stimulus, the game engine was used to compute a ground-truth semantic segmentation of the video. Brain activity was recorded with fMRI in both experiments. Voxelwise encoding models were then fit separately for the subjects in the two experiments to capture the semantic tuning of the brain during navigation and movie-watching. The visual semantic tuning of the brain in the two experiments was then compared to determine the visual semantic tuning differences between movie-watching and navigation.

## MATERIALS AND METHODS

In this study we compared visual semantic tuning in the human cerebral cortex in data collected from two experiments. Participants in the first experiment watched movies, while participants in the second experiment engaged in active navigation. Voxelwise modelling was used to recover visual semantic tuning in both experiments, for every cortical voxel in every participant. Because the data from the movie-watching and navigation experiments were collected from different participants and exist in different anatomical spaces, freesurfer was used to map data across anatomical spaces of the participants.

### Participants

Five participants (4 male, 1 female, ages 25-32) participated in the movie-watching experiment and six participants (3 male, 3 female, ages 24-34) participated in the driving experiment. All participants were healthy adults with normal or corrected to normal vision. The experimental procedures were approved by the Institutional Review Board at the University of California, Berkeley, and written informed consent was obtained from all participants.

### Movie-watching experiment

#### Task

The movie data was collected following the procedures outlined in (Nishimoto et al., 2011) and (Huth et al., 2012). Briefly, natural movies were drawn from the Apple QuickTime HD gallery and Youtube, and cropped to 512 x 512 pixels. Multiple 10-20 second clips were concatenated to form 21 10-minute runs. 12 runs were used as model training runs, and 9 runs were used for model validation. Five male participants (ages 25-32) watched these movies at fixation while BOLD activity was recorded.

#### Feature spaces

To create visual semantic features, an observer manually tagged every second of the movies with WordNet labels (Miller, 1995). Categories were tagged if they appeared in at least half of each second. One-hot encoding features were then constructed for all categories that appeared in the movies. Additionally, features were also included for all superordinate categories for the labelled categories. For example, in addition to “dog,” features were also included for “canine” and “domestic animal.” There were 1,364 object categories and 341 superordinate categories, for a total of 1,705 features.

Because the high-level semantic content and the low-level visual structure of naturalistic movies are correlated, models fit to only the visual semantic features may be confounded by stimulus correlations between high- and low-level features. To account for these stimulus correlations in subsequent analyses, motion-energy features (Nishimoto and Gallant, 2011) were also computed for these movies to capture their low-level visual structure. To create this feature space, the screen was tiled with 6,555 spatiotemporal quadrature Gabor filters of varying size, orientation, and temporal frequency, and the output of the filters applied to the movie were used as features.

### Navigation experiment

#### Task

The navigation data was collected in (Zhang et al., 2025). Briefly, participants performed a taxi-driver task in a virtual city while brain activity was recorded with fMRI. A large dynamic virtual city was constructed with Unreal Engine 4 and CARLA (Dosovitskiy et al., 2017), and was populated with both vehicular and pedestrian traffic. Participants used a custom MR-compatible steering wheel and pedal set to drive a car through their virtual world from a first-person perspective. Prior to scanning, participants learned the layout of the world. In the scanner, the taxi-driver task was divided into trials. On each trial, a participant was cued to navigate to a particular location in the world. The participants were instructed to use their knowledge of the world to drive to the cued location via the quickest path, while respecting all traffic rules. After the participant arrived at the destination, a new trial began. Participants were allowed to freely view the screen while driving, and eyetracking data was collected at 60 Hz. Trials were recorded with Unreal Engine’s built-in demo recording system. Data was collected in 11-minute runs. 120 minutes of data was collected from participant 1, and 180 minutes of data each was collected from participants 2-6.

#### Feature spaces

To compute visual semantic features, the game engine was used to semantically segment the participant’s view to determine the category of the object rendered at each pixel. This semantic segmentation covered 16 categories that were relevant to driving: buildings, pedestrians, vehicles, roads, road lines, sidewalks, traffic signs, poles, walls, fences, fields, ground, foliage, self, miscellaneous objects, and sky. Semantic categories were also included for on-screen display items that were overlaid on the screen rather than rendered in-world: the “go to” and “arrived” cues and the speedometer. The semantic feature space also included 14 additional categories for objects and on-screen display items that appeared during the learning phase, but not during data collection.

Because participants are unlikely to allocate equal attention to all areas of the screen, the semantics of the entire screen may not accurately reflect representations of visual semantics with the influence of top-down attention. Therefore, the gaze location was used as a proxy of how participants allocated attention. For each sample in the eyetracking data, a circle of 5° diameter was drawn around the gaze position on the corresponding semantic segmentation frame. Then the fraction of this circle that was occupied by each of the 16 categories was calculated and used as features.

In the navigation experiment, the visual semantic content of the scene co-varies with many other features, such as the low-level visual structure of the scene, the participants’ motor actions, and other navigation-related parameters. The contribution of the representation of these other features to brain activity were captured by 37 additional feature spaces that are not described here (but see (Zhang et al., 2025)). In total, these 37 feature spaces encompassed 28,101 additional features.

### Voxelwise encoding models

Voxelwise modeling (Naselaris et al., 2011; Nishimoto et al., 2011; Huth et al., 2012, 2016; Deniz et al., 2019; Lescroart and Gallant, 2019; Nunez-Elizalde et al., 2019) was used to quantify semantic tuning across the brain in each of the two experiments. VM models the time series of the BOLD activity in each voxel as a linear combination of the time series of all features from all feature spaces. In the movie-watching experiment, we fit models for the 1705 Wordnet visual semantic features and 6555 motion-energy features to the BOLD activity of each voxel. In the navigation experiment, we fit models for the 16 visual semantic features along with 28,101 other features in 37 additional feature spaces. To compensate for differences in the size of feature spaces and for correlations between features, banded ridge regression was used to estimate the optimal model weights (Nunez-Elizalde et al., 2019; Dupré la Tour et al., 2022). To capture the hemodynamic response, an FIR filter was implemented by including delayed copies of all features at 5 temporal delays. The optimal model weights and shape of the FIR filter for each voxel were empirically selected by cross-validation. Model performance was then quantified by the coefficient of determination (R^2^) between the predicted and actual brain activity on a hold-out dataset not used during model training. Permutation tests were used to estimate statistical significance in each voxel. A significance mask was determined at an FDR corrected p < 0.05 level in each participant. For each voxel, the joint model performance for all models was then partitioned over models for each of the feature spaces (2 in the movie-watching experiment, and 38 in the navigation experiment). Here we call these values the split R^2^ scores. This study focuses on the visual semantic models fit in the two experiments (For visual semantic model performances see Supplemental Figures 1 and 2).

**Figure 2.**
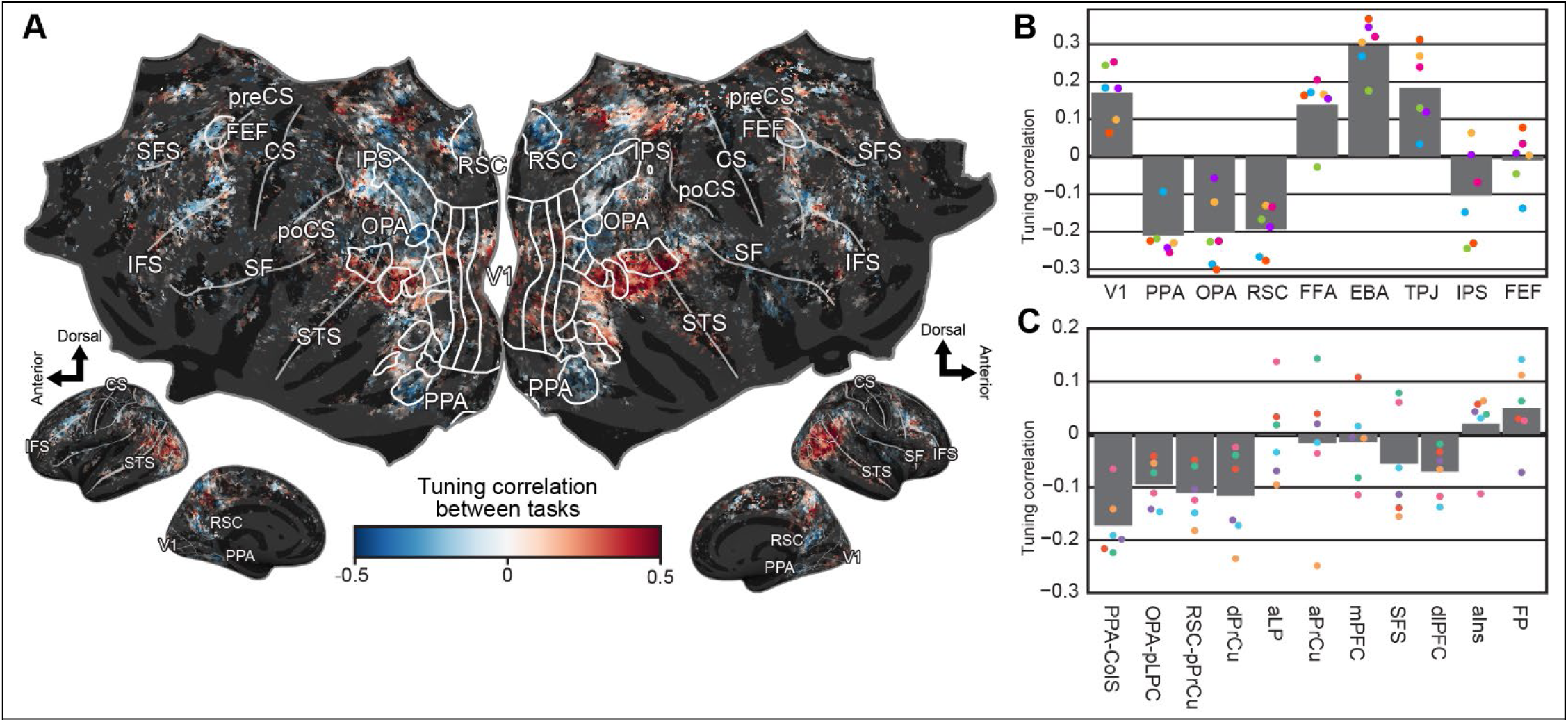
Visual semantic tuning across the cortex shifts between movie-watching and navigation. Different task demands may cause the brain to shift its visual semantic tuning between movie-watching and navigation. A) The magnitude of the tuning shift was quantified as the correlation between the visual semantic tunings in the two experiments at each vertex on the brain. A higher correlation corresponds to a weaker tuning shift, and a lower correlation to a stronger tuning shift. The tuning in each navigation participant was compared with the mean tuning in the movie-watching participants. Results show that visual semantic tuning shifts significantly between tasks in 42.0 ± 6.3% of the cortex (mean ± stdev across driving participants, p < 0.05, FDR corrected bootstrap test). Significant tuning shifts are shown on the group-level map. For correlations in individual navigation participants to the movie-watching mean, see Supplemental figure 3. B) Because movie-watching and navigation likely engage different functional networks, the visual semantic tuning shifts may differ across functional regions. The average tuning correlation between the two tasks are shown for selected ROIs. Individual navigation participants are shown in colored dots. The tuning shifts are the strongest in PPA, OPA, and RSC, three scene-selective regions, followed by IPS and FEF, two visual attention regions. The tuning shifts are the weakest FFA, EBA, and TPJ, which are people- and socially-selective, which are comparable to the tuning shift in V1. These results suggest that while human-selective regions have more consistent visual semantic tuning across tasks, visual attention regions and place-selective regions are more flexible and adapt more to task demands. C) In other work we had shown the functional network mediating active navigation is also selective for visual semantics. Here, we examined how visual semantic tuning in these regions differed between movie-watching and navigation. Bars show the group-average correlation in visual semantic tuning between the experiments in each region, and dots indicate individual subjects. All but two regions show a zero or negative correlation in visual semantic tuning between experiments, suggesting that the navigation network flexibly shifts its visual semantic representations across tasks. These results show that the spatial navigation network, like visual attention regions, flexibly shifts its visual semantic tunings when engaged during navigation.

To recover the visual semantic tuning of each voxel in each experiment, the fit model weights were averaged across the five delays. However, because banded ridge regression optimizes the regularization parameter for each feature space in each voxel, the scale of the model weights differs across voxels. To account for this scaling difference, the recovered vectors were first normalized to have unit norm, and then scaled by the split R^2^ score of the visual semantic model. This rescaling effectively corrects for the differences in the scales of model weights caused by different regularization hyperparameters.

### Localizers for known functional ROIs

Separate functional localizer experiments were run to identify conventional functional ROIs in each participant. ROIs were determined using five sets of localizers: a retinotopic localizer, an MT localizer, a visual category localizer, a motor localizer, and a theory-of-mind localizer.

#### Retinotopic Localizer

Retinotopic localizers were collected in four 9-minute runs. In two runs, rotating wedges were used to determine visual angles. In the other two runs, expanding rings were used to determine eccentricity. Visual angle and eccentricity maps were then used to delineate V1, V2, V3, V4, V3A, V3B, and V7 (Hansen et al., 2007).

#### MT Localizer

MT localizer data were collected in four 90-second runs. These runs consisted of alternating 16-second blocks of motion coherent and incoherent random dot fields. The contrast between coherent and incoherent blocks was used to delineate MT+.

#### Visual category localizer

Visual category localizer data were collected in six 4.5-minute runs. These runs consisted of 16 blocks of 16 seconds each. In each block, 20 images of either places, faces, body parts, non-human animals, household objects, or phase-scrambled objects were displayed. Each image was displayed for 300 ms followed by a 500 ms blank. The contrast between faces and objects was used to define the fusiform face area (FFA, (Kanwisher et al., 1997)). The contrast between body parts and objects was used to define the extrastriate body area (EBA, (Downing et al., 2001)). The contrast between places and objects was used to define the parahippocampal place area (PPA, (Epstein and Kanwisher, 1998)), occipital place area (OPA (Dilks et al., 2013)), and retrosplenial cortex (RSC, (Aguirre et al., 1998)).

#### Motor localizer

Motor localizer data were collected in one 10-minute run. Participants were cued to randomly alternate between six motor tasks in 20-second blocks. For “hand,” “foot,” “mouth,” “speech,” and “rest,” the cue was simply that word presented at the center of the screen. For the saccade condition, participants were shown a screen filled with randomly distributed saccade targets.

In the “hand” blocks, participants were instructed to make small finger-drumming movements. In the “foot” blocks, participants were instructed to make small toe movements. In the “mouth” blocks, participants were instructed to make small lip movements without moving the tongue or jaw. In the “speak” blocks, participants were instructed to continuously subvocalize self-generated sentences without physically speaking. In the “saccade” blocks, participants were instructed to continuously saccade between targets presented on the screen. A linear model was used to find the change in BOLD response in each voxel in each condition relative to the mean BOLD response.

Weights for the hand, foot, and mouth responses were used to delineate the primary motor and somatosensory areas for hands (M1H, S1H), feet (M1F, S1F), and mouth (M1M, S1M). Weights for the saccade responses were used to delineate the intraparietal sulcus (IPS), frontal eye fields (FEF, (Paus, 1996)), and frontal operculum (FO, (Corbetta et al., 1998)). Weights for the “speak” responses were used to delineate Broca’s area and the superior ventral premotor speech area (sPMv).

#### Theory-of-mind localizer

Theory of mind localizer data were collected in two 5-minute runs. Participants were shown a series of short stories consisting of false belief, false photograph, desire, physical description, and nonhuman description stories (Saxe and Kanwisher, 2003). Each story was shown for 10 seconds. After each story, participants were presented with a fill-in-the-blank two-alternative forced choice question for 4 seconds and responded with a buttonbox. Half of the questions concerned the belief of characters in the stories, while the other half were concerned with factual information. The contrast between belief and factual questions was used to delineate the temporoparietal junction (TPJ).

### Comparison of semantic tuning across tasks

In order to compare visual semantic tuning in the cerebral cortex between participants in the movie-watching and navigation experiments, we must map between the anatomical spaces of the participants and also between the visual semantic feature spaces used by the two experiments.

#### Mapping between participants

The data from the movie-watching and navigation experiments were collected from different participants. To compare visual semantic tuning between participants from the two experiments, the data must first be in the same anatomical space. Because visual semantic tuning is largely consistent across participants (Huth et al., 2012; Visconti di Oleggio Castello et al., 2025), the simplest approach would be to compare group-average models from both experiments in a template space. Creating a group-average model is well-suited for data from the movie-watching experiment, which uses a purely stimulus-driven task with no behavioral variability. However, because the navigation task involves active behavior that varies across individuals, a group-average model is less well-suited for this experiment. To preserve individual differences in navigation, we therefore mapped the group-averaged movie-watching data to each individual participant in the navigation experiment, and compared each navigation participant with the group-average move-watching data.

To map between anatomical spaces, we first mapped the cortical voxels recorded in each moving-watching participant into the fsaverage6 atlas space. This mapping was implemented in several steps. First, the surf2surf function in freesurfer (Dale et al., 1999) was used to calculate a mapping from the native space of each movie-watching participant to the fsaverage surface. This mapping was used to project semantic tuning vectors from each movie-watching participant to the fsaverage surface. Then, surf2surf was used to compute a mapping from the fsaverage surface to each navigation participant’s surface. The group average movie-watching visual semantic tuning map was then mapped from fsaverage to the cortical surface of each navigation participant. Finally, the visual semantic tuning between movie-watching and navigation was compared at each vertex in the native cortical space of each navigation participant.

#### Comparing across visual semantic feature spaces

The visual semantic feature space created for the movie-watching experiment consisted of 1705 semantic features drawn from common WordNet categories. In contrast, the visual semantic feature space created for the navigation experiment consisted of 16 distinct features defined by the semantic segmentation component in the game engine. Therefore, to assess tuning shifts between experiments, these feature spaces needed to be aligned by defining a shared semantic subspace. Although the WordNet feature space covers a large number of semantic categories, it did not encompass all 16 semantic categories in the navigation experiment. We excluded four visual semantic categories in the navigation experiment that did not correspond to WordNet categories: “None” labeled untagged objects, “other” served as a catch-all for unlabeled objects, “road lines” represented a driving-specific category absent from movies, and “self” did not appear due to the third-person perspective of movie stimuli. We also excluded “walls” because participants encountered no wall-labeled objects in the virtual environment, preventing tuning estimation despite its presence in both taxonomies. The shared semantic space thus spanned 11 categories: buildings, fences, people, poles, roads, sidewalks, foliage, vehicles, traffic signs, fields, and ground.

Because of the hierarchical nature of WordNet, there was a many-to-one correspondence between the WordNet semantic features in the movie-watching experiment and the disjoint semantic segmentation features in the driving experiment. For example, the category “Buildings” includes many subordinate categories such as “skyscraper,” “barn,” and “house,” among others. To account for this many-to-one correspondence, the WordNet visual semantic tuning was averaged across each category with all of its subordinate categories. Doing so produced 11 non-hierarchical categories directly comparable to the semantic categories in the navigation experiment.

### Quantifying the semantic tuning difference between tasks

Once the movie-watching and navigation tuning were anatomically aligned, the tuning differences between the experiments could be evaluated. In prior work we used a tuning shift index (TSI) to quantify such differences (Çukur et al., 2013). However, the TSI measures the strength of tuning shifts along a known axis in semantic space. Because in this study we have no *a priori* knowledge of what specific semantic categories were attended in either experiment, the TSI is not an appropriate measure. Instead, we quantified both the direction and magnitude of task-related tuning shifts in semantic space. The strength of the tuning shift was quantified as the correlation between the two tuning vectors at each vertex. A correlation of 1 indicates that there is no semantic tuning shift, a correlation of -1 indicates that the vertex shifts to opposite tunings between the two experiments, and a correlation of 0 indicates that the vertex shifts to orthogonal tunings between these two experiments. The direction of the semantic tuning shift was quantified as the difference between the tuning vectors in the two experiments. To account for the difference in weight vector scales between the experiments, the norms of weight vectors in each participant were first separately normalized across the cortex such that the 2^nd^ percentile became 0 and the 98^th^ percentile became 1, and then the normalized movie-watching weight vector at each vertex was subtracted from the navigation weight vector. The resulting vector describes the tuning shift direction at each vertex.

### Bootstrap testing of the significance of tuning differences

A bootstrap procedure was used to test whether any visual semantic tuning differences between movie-watching and navigation were statistically significant at each vertex in each navigation participant. If visual semantic tuning at each vertex remained constant across tasks, the correlation between visual semantic weights from movie-watching and navigation should be close to 1. If visual semantic tuning differed across movie-watching and navigation, then the correlation should be significantly less than 1. A bootstrap procedure was used to estimate the distribution of these correlations. To establish a null distribution describing the expected correlation values for when no tuning shifts occur, the movie-watching semantic weights were resampled 1,000 times. On each bootstrap, the data was resampled in 20-TR blocks of the feature and BOLD timeseries, and the encoding model was refit to this resampled data. The weight from these refit models were then correlated against the original movie-watching weights to form a null distribution for when no tuning shift occurs. The navigation semantic weights were then also bootstrapped 1,000 times using the same procedure. On each iteration, bootstrapped navigation semantic weights were correlated with the movie-watching semantic weights to obtain a distribution of between-task correlations. Statistical significance was determined at each vertex by comparing these two distributions.

### Comparing the representation of object categories between tasks

The analyses above characterized the tuning of individual cortical vertices to different object categories. The aggregate effects of tuning shifts across the cortex may also cause the spatial organization of functional networks to change across tasks. Therefore, we also examined how task context affects the cortical distribution of visual semantic tuning for each category between the movie-watching and navigation experiments.

The difference in the representation of object categories between the movie-watching and navigation tasks were quantified as the spatial correlation across the cortex for the weight maps for each of the object categories between the two tasks. A correlation of 1 indicates that the object category is represented by the same brain regions, while a correlation of -1 indicates that object category is represented by the inverse of set of functional regions between the two experiments, and a correlation of 0 indicates that the object category is represented by orthogonal sets of brain regions these two experiments. The same bootstrap procedure described above to assess tuning differences was used to determine whether the representation of object categories was significantly different between movie-watching and navigation. On each of the 1,000 bootstraps of the movie-watching weights, we correlated the bootstrapped weights for each object category with the original weights to obtain a distribution of the expected null correlations. Statistical significance was determined for each object category by comparing the actual correlation with this bootstrapped null distribution.

### Animacy Index

Because previous experiments had identified the contrast between animate and inanimate objects as a primary organizational principle for object representations (Naselaris et al., 2012; Grill-Spector and Weiner, 2014), we hypothesized that the cortex might reorganize object representations along this animacy dimension to reflect their behavioral relevance between tasks. To test this hypothesis, we defined an *animacy index* to quantify where the representation of each object category fell along the animate-inanimate spectrum. We first used an earlier study (Naselaris et al., 2012) to identify a list of animate and inanimate objects. Using the movie-watching weights as a reference baseline, we computed template “animate” and “inanimate” representations by averaging the visual semantic weights separately across all animate categories and across all inanimate categories. The *animacy index* for any object category was then computed as the difference between its correlation with the animate template and its correlation with the inanimate template, normalized by the correlation between the two templates.

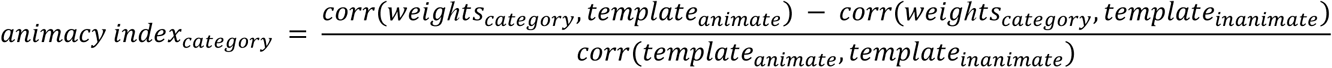

This animacy index ranges from -1 (fully inanimate representation) to +1 (fully animate representation). The animacy index was then computed for each of the 11 shared semantic categories separately in each participant in the movie-watching and navigation experiments. To determine whether the animacy index of an object category differs between tasks, we used a permutation test. Because each participant constitutes one sample for the representation of each object category, permutation of group membership participants across movie-watching and navigation were used to determine a null distribution for difference in animacy indices.

### Code availability

Supporting code will be made available on github upon publication.

## RESULTS

Prior studies suggest that attention improves task performance by dynamically increasing the representation of task-relevant information in the brain, while decreasing the representation of irrelevant information. However, those studies only examined these effects under a narrow range of highly controlled conditions. Few prior studies have sought to examine how attention might change the representation of information across complex, naturalistic tasks (Kay et al., 2023).

Movie-watching and navigation are two naturalistic tasks that likely require participants to attend to different visual semantic categories in order to perform each task optimally. Therefore, to examine how attention affects the representation of task-relevant information under naturalistic conditions, we compared visual semantic tuning across the cerebral cortex in a group of participants who passively watched narrative movies (Huth et al., 2012) versus visual semantic tuning in a different group of participants who navigated through a virtual world (Zhang et al., 2025). BOLD activity was recorded with fMRI in both experiments, and voxelwise encoding models were used to capture visual semantic representations in the brain in the two experiments separately.

### Visual semantic tuning differs between movie-watching and navigation

To determine whether visual semantic representations and tuning differ between the two tasks, we compared visual semantic weights obtained from the participants who watched movies to those obtained from participants who navigated in a virtual world. The average visual semantic weights from the movie-watching participants were mapped to the cortical surface of each navigation participant (see methods). For each navigation participant, the visual semantic weights during navigation at each cortical vertex was compared with the group-average movie-watching weights projected into that participant’s anatomical space. Here, a correlation that is significantly less than 1 indicates that visual semantic tuning differed between the two tasks. Results show that visual semantic weight vectors differed significantly across much of the cortex between the two tasks. Between movie-watching and navigation, the correlation was significantly less than 1 in 42.0 ± 6.3% of cortical vertices (mean ± stdev across navigation participants, p < 0.05, FDR corrected bootstrap test, Figure 2A). The correlation between visual semantic tunings in the two tasks is shown on the flattened group-level cortical surface in Figure 2A (for comparisons of individual driving participants with the movie-watching participants, see Supplemental Figure 3). These results show that there are widespread cortical visual semantic tuning shifts across tasks.

Because movie-watching and navigation likely engage different functional networks, we sought to determine whether the visual semantic tuning shifts differ across functional regions. Inspection of Fig 2A suggests that the strength of the tuning difference differs systematically across brain regions. To better understand region-specific tuning differences, we determined the average correlation within regions of interest (ROIs) that are known to be attention-related or selective for visual semantic categories (Fig. 2B). Results show that tuning shifts are the strongest in the parahippocampal place area (PPA), occipital place area (OPA), and retrosplenial cortex (RSC), followed by the intraparietal sulcus (IPS) and frontal eye fields (FEF); tuning differences are weakest in the fusiform face area (FFA), extrastriate body area (EBA), temporal-parietal junction (TPJ), and the primary visual cortex (V1). In sum, the weakest shifts were found in human-related and social ROIs (FFA, EBA, and TPJ), and the strongest tuning shifts were found in place-selective (PPA, OPA, and RSC) and visual attention (IPS and FEF) ROIs. These results suggest that while human-selective regions show the most consistent visual semantic tuning between movie-watching and navigation, place-selective and visual attention regions are more flexible and change their visual semantic tunings to adapt to the different demands of the two tasks.

### Tuning differences are not related to differences in stimulus statistics

One potentially important confound in this study is that the visual semantic stimulus statistics differed between the movie watching and navigation experiments. For example, “people” occurred much more frequently in the movie-watching experiment than in the navigation experiment. It is theoretically possible that the observed tuning differences might be an artifact of these differences in stimulus statistics. We used two convergent methods to test that stimulus statistics did not confound our results. First, we computed the relative differences between the two experiments across each of the 11 visual semantic categories evaluated in this study (Supplemental Figure 4). The amount of variance in the observed tuning shifts that could be explained by stimulus differences between movie watching and navigation is not significant (11.4 ± 2.8 % mean ± stdev across participants, chance = 9.9%, p > 0.05, permutation test). In a second test we computed the correlation between the stimulus differences and the principal components of the tuning shifts. The stimulus statistic differences were not significantly correlated with any of the PCs of the tuning shifts (p > 0.05, permutation test). Taken together, these two analyses suggest that the observed semantic tuning shift between movie-watching and navigation are intrinsic to the brain, and not a mere artifact of the difference in stimulus statistics in the two experiments.

### Tuning shifts differ across functional regions

While the analyses above show that visual semantic tuning differed between movie-watching and navigation, they do not reveal which specific semantic categories the brain prioritizes in each task. To characterize semantic targets of the tuning shifts between movie-watching and navigation, we subtracted the average movie-watching weights from the navigation weights at each vertex in every navigation participant, and then applied principal components analysis (PCA) to these tuning differences. Across all six navigation participants, we find three PCs that explain significant amounts of variance in these tuning differences (p < 0.05, permutation test), accounting for 53.9 ± 6.69 % of the total variance (mean ± stdev across driving participants, Supplemental Figure 5).

Figure 3A shows these three PCs on the group-level cortical surface, mapped to the red, green, and blue color channels respectively (for individual navigation participants see Supplemental Figure 6). Figures 3B and 3C show visual semantic tuning during movie-watching and navigation on these same PCs. Inspection of Figure 3 reveals, the IPS and FEF, two well-known regions that mediate visual attention, shift their visual semantic tuning towards sidewalks, traffic signs, and buildings during navigation compared to movie-watching. The PPA and RSC, two place-selective regions, and the inferior frontal sulcus shift their visual semantic tuning towards ground and fields. Many other cortical regions, including precuneus, TPJ, insula, and prefrontal cortex, shift their tuning towards vehicles. (Average tuning shifts in individual ROIs are shown in Supplemental Figure 7). These results suggest that functional regions separately reconfigure their visual semantic representations between tasks, likely reflecting complex interactions between their baseline representations and their task-specific roles.

**Figure 3.**
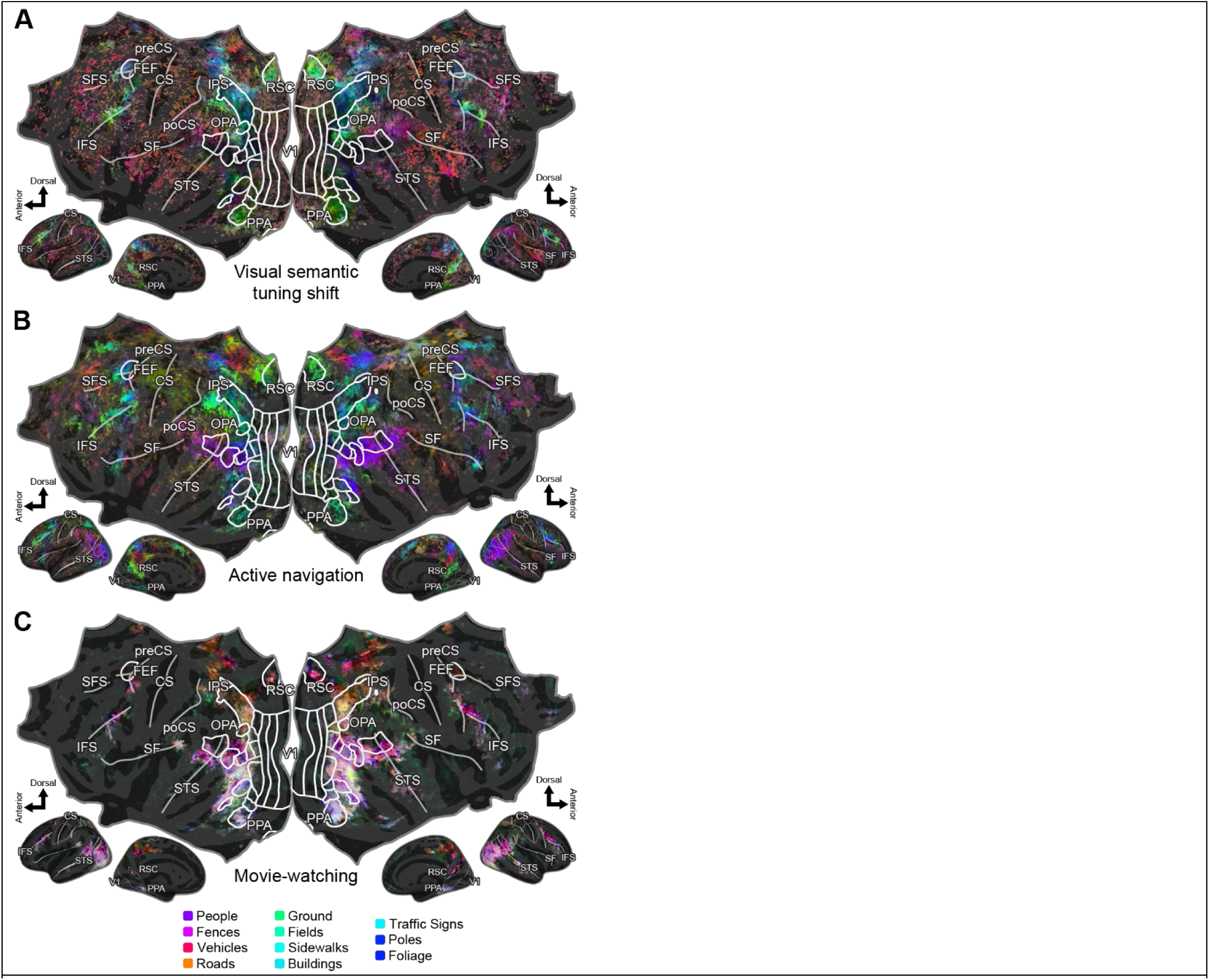
Visual semantic tuning shifts vary across the cortex. While the correlation analysis shows that visual semantic tuning differed between the two tasks, it does not reveal how the cortex reorganizes its selectivity across semantic categories. To identify the specific differences in semantic selectivity between movie-watching and navigation, we subtracted the visual semantic weights during movie-watching from the visual semantic weights during navigation at each vertex on the brain. Principal components analysis identified three significant group-level PCs in these 11-dimensional tuning shift vectors (p < 0.05, permutation test). To visualize these tuning shifts, we projected the tuning shifts on to these three PCs, which are mapped to the red, green, and blue color channels respectively. To aid in interpretation, the 11 visual semantic categories were also projected into this space, and their colors are shown at the bottom. A) Visual semantic tuning shifts between active navigation and movie-watching are shown on the group level map (for individual navigation participants see Supplemental Figure 6). The IPS and FEF show a shift towards sidewalks, traffic signs, and buildings. The PPA, RSC, and inferior frontal sulcus shows a shift towards ground and fields. The rest of the cortex, including precuneus, TPJ, insula, and prefrontal cortex, show a shift towards vehicles. B) and C) For comparison, the visual semantic tuning in navigation and movie watching were also projected into this space. These maps reveal a striking difference in visual semantic tuning across the cortex between tasks, and show that tuning shifts vary systematically across ROIs. These results suggest that between tasks, the brain shifts its visual semantic tuning in a complex manner to optimize the use of limited cortical resources.

### Navigation-related regions show strong tuning shifts

In an earlier study we reported that human navigation is supported by a cortical network of 11 different functionally distinct areas distributed across visual, parietal and prefrontal regions of the brain (Zhang et al., 2025). This network represents a wide variety of perceptual, cognitive, and motor information directly relevant for navigation. If attention shifts tuning to increase the representation of task-relevant information, then we should expect profound changes in visual semantic tuning between passive movie-watching and navigation will occur in the functional areas comprising the navigation network. We find that these 11 regions showed relatively strong differences in tuning between movie-watching and navigation (Figure 2C). Nine regions (occipital areas PPA-ColS, OPA-pLPC, and RSC-pPrCu; parietal areas aLPC, aPrCu, and dPrCu; and prefrontal areas dlPFC, mPFC, and SFS) shift their visual semantic tuning toward landmarks such as sidewalks, buildings, and fields, and away from vehicles and pedestrians. (The precise pattern of visual semantic tuning shifts varies across these areas, see Supplemental Figure 8 for details.) However, two areas within the navigation network (prefrontal areas aIns and FP) do not show a clear tuning shift direction. In sum, these results confirm the hypothesis that attention to task-relevant information changes visual semantic representations broadly across task-relevant functional brain regions in order to optimize task performance.

### Tuning shifts enhance navigation-relevant information

Tuning shifts serve to increase representation of task-relevant information. To identify the primary semantic dimensions the brain prioritizes during navigation compared to movie-watching, we examined the loading of the 11 categories on the PCs of the tuning shifts (Figure 4). PC 1 is an almost singular shift towards vehicles, suggesting the brain prioritizes the representation of other agents. PC 2 shows a shift towards buildings, ground, fields, foliage, and traffic signs, and a shift away from vehicles, roads, sidewalks, and people, suggesting that the brain seeks to better discriminate between scene identity-relevant categories and the entities that populate the scene. PC 3 shows a shift primarily towards people and away from roads, suggesting that the brain also seeks to better discriminate between navigational affordances and people. These results show that during navigation, the brain shifts its visual semantic selectivity to better identify other agents in the environment and to discriminate these agents from navigational cues.

**Figure 4.**
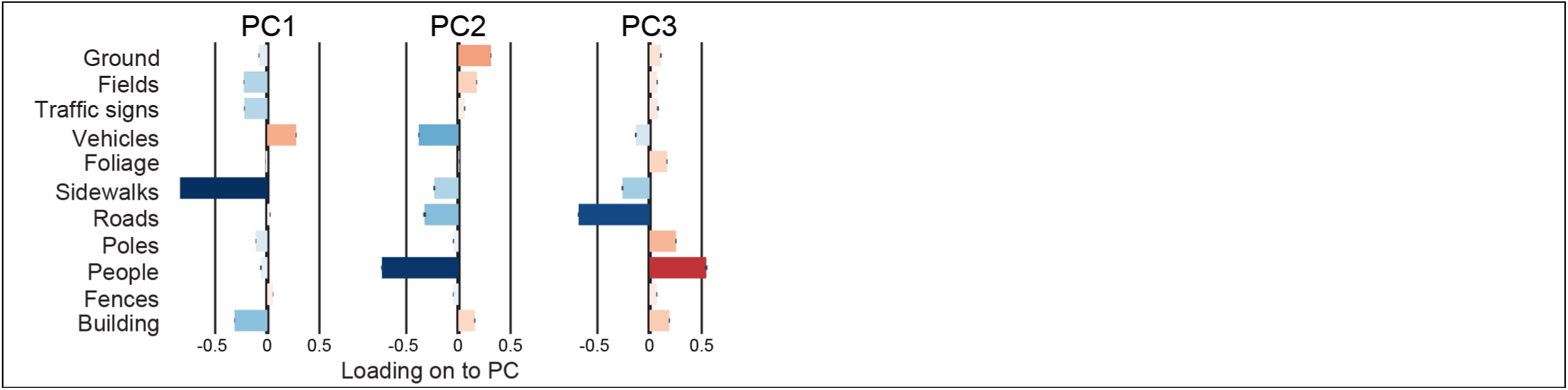
Loadings of the 11 visual semantic categories onto the three group-level principal components. Tuning shifts serve to increase the representation of task-relevant information across the brain. To understand the semantic dimensions that the brain re-prioritizes between navigation than movie-watching, we examined the loadings of the 11 categories on to the three significant group-level principal components (PCs) of the tuning shifts. PC 1 shows an almost singular shift towards vehicles during navigation, and away from sidewalks, buildings, traffic signs, and fields, suggesting that the primary tuning shift enables the brain to better identify other vehicles in the environment. PC 2 shows a shift towards buildings, ground, fields, foliage, and traffic signs, and away from vehicles, roads, sidewalks, and pedestrians, suggesting this shift enables the brain to better discriminate between scene identity-relevant categories and the entities that populate the scene. PC 3 shows a shift away from roads, sidewalks, and vehicles, and primarily towards pedestrians and away from roads, suggesting that this shift enables the brain to better discriminate between navigational affordances and pedestrians. These PCs suggest that the brain shifts its visual semantic representations to better represent and differentiate between task-relevant object categories.

### Tuning shifts change the representation of object categories

These tuning shifts show that cortical regions change their semantic selectivity between tasks. A possible effect is that each object category may therefore be represented by different functional networks across tasks. To test this hypothesis, we examined the spatial correlation of weights for each object category across the cortical surface between the movie-watching and navigation in each navigation participant. The representation of buildings, people, poles, roads, sidewalks, foliage, and vehicles were significantly different between movie-watching and navigation (p < 0.05, FDR corrected, bootstrap test, Supplemental Figure 9). The difference in representation is not constant across categories: the representation of people is most correlated across tasks (r = 0.259 ± 0.075 mean ± stdev across navigation participants) while the representation of vehicles is least correlated (r = -0.078 ± 0.085 mean ± stdev across navigation participants). This reorganization suggests that across tasks, the brain redistributes object category representations across functional regions, with task-critical categories, e.g. vehicles, undergoing the biggest shift.

While direct comparison of weight vectors reveals that the brain systematically reorganizes object representations between tasks, it remains unclear whether these differences follow interpretable organizational principles. The animate-inanimate distinction is a primary organizing principle of object representations in the cortex (Naselaris et al., 2012; Grill-Spector and Weiner, 2014), and the perceived animacy of even simple geometric shapes can be affected by context such as their motion (Heider and Simmel, 1944). Thus, we hypothesized that the cortex might reorganize object representations along this animacy dimension to reflect their behavioral relevance between tasks. Specifically, objects that appear to move intentionally might be represented in a more animate manner during navigation, even if they are typically considered inanimate objects. To test this hypothesis, we defined an *animacy index* for the representation of object categories in each movie-watching and navigation participant. Results show that the animacy of roads, vehicles, traffic signs, and the ground differed significantly between the two experiments (FDR corrected p < 0.05, permutation test, Fig 5A, B). The representation of vehicles showed the largest difference: vehicles are represented as inanimate objects during movie-watching, but animate objects during navigation (FDR corrected p < 5e-3, permutation test). Roads and traffic signs representations also are more animate during navigation, while ground representation is more inanimate during navigation. To visualize these differences intuitively, object category representations were mapped onto the flattened cortical surface. Figure 5C shows the representation of vehicles, the category with the strongest animacy difference between tasks, while Figure 5D shows the representation of people, a category whose animacy did not change between tasks (for other categories, see Supplemental Figure 10). Vehicle representations occupy distinct cortical networks between navigation and movie watching, while people representations occupy similar networks across tasks. These results demonstrate that the brain flexibly reconfigures the perceived animacy of objects across task contexts.

**Figure 5.**
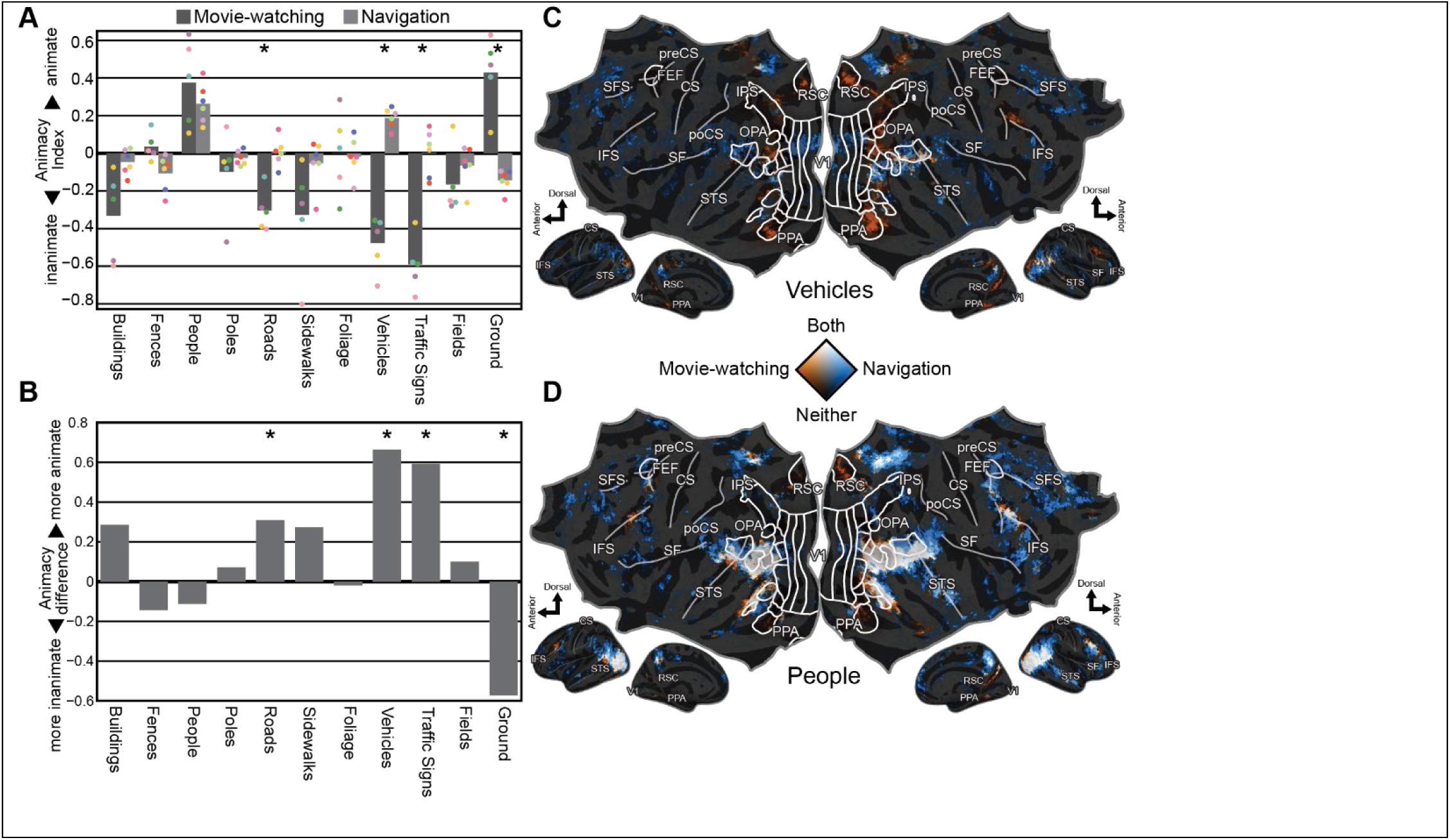
The animacy of object category representations differs between tasks. Animacy is a primary organizational principle for object category representations in the cortex. Because tuning shifts cause object categories to be represented by different cortical regions, this shift may correspond to changes in the animacy of object category representations. To quantify this, we defined an animacy index that quantified how object category representations project to template “animate” and “inanimate” representations. The animacy indices for the representation of the 11 object categories are shown (bars indicate group average values while dot indicate individual navigation participants). The animacy of the representation of roads, vehicles, traffic signs, and the ground are different between the two tasks at the group level (FDR corrected p < 0.05, permutation test). Notably, the representation of vehicles is inanimate during movie-watching, while animate during navigation. The representations for roads and traffic signs are less inanimate in navigation than in movie-watching, while the representation of ground shifts from animate to inanimate (see discussion on the change in the representation of “ground”). These results show that one effect of task-related tuning shifts is a difference in the animacy of object category representations, and that this difference is specific to task-relevant categories.

## DISCUSSION

In this study, we hypothesized that the human brain shifts its visual semantic tuning between naturalistic tasks. To test this hypothesis, we compared visual semantic tuning measured from participants who passively watched movies and from participants who actively navigated in virtual reality. Results show that visual semantic tuning across the cerebral cortex differs significantly between the two tasks. The magnitude of the shifts in visual semantic tuning varies across ROIs; the strongest shifts are found in regions that represent information about navigation and spatial attention. These results show that when people switch from one task to another, many visually-selective regions of the brain shift their tuning.

Post-hoc inspection suggests that these visual semantic tuning shifts are directly related to task demands. During active navigation, the precuneus, temporoparietal junction (TPJ), and prefrontal cortex shift tuning to increase the representation of vehicles. These regions have been implicated in theory-of-mind and higher-level planning (Koechlin et al., 2000; Saxe and Kanwisher, 2003; Kaller et al., 2011). Therefore, this pattern likely reflects the greater behavioral importance of vehicles during navigation, where they directly influence participants’ driving decisions, compared to movie watching, where they serve as background objects. The engagement of TPJ, a theory-of-mind region, suggests that the enhanced vehicle representation may support cognitive processes involved in predicting the future actions of other agents. Active navigation also increases the representation of traffic signs and poles. However, this tuning shift is localized to visual scene perception areas including RSC, OPA, and PPA, as well as spatial attention regions including the IPS and FEF. Thus, this pattern likely reflects the greater behavioral importance of traffic signs during navigation, where they govern permissible actions, compared to movie watching, where they serve as background props. The cortical networks showing increased representation of vehicles and traffic signs are distinct, suggesting that representing other agents and representing spatial rules engage different cognitive mechanisms.

The difference in the representation of vehicles between movie watching and navigation is striking: in movie-watching, vehicles are represented as inanimate objects, while in navigation, they are represented as animate objects. While it has long been known that interactions of simple moving geometric objects can influence their perceived animacy (Heider and Simmel, 1944; Santos et al., 2008, 2010; Osaka et al., 2012; Schultz and Bülthoff, 2019), previous works have studied animacy only in the context of abstract geometric motion. Here, we compared the animacy of real-world object categories across naturalistic tasks, and find that the animacy of the same object category differs across task contexts. This effect cannot be attributed to visual co-occurrence of drivers with vehicles, because the car models in the virtual environment did not include visible drivers. Instead, task context likely drove the representational shift. During movie watching, vehicles serve as background objects or human-operated conveyances, whereas during navigation, participants embody a virtual car and other vehicles become peer agents sharing the environment. The agency of other vehicles during navigation is reflected in their animate cortical representation. Unexpectedly, roads and traffic signs were also represented more similarly to animate objects during navigation despite not being objects with agency. This effect may reflect that roads and traffic signs, while not animate, directly influence participants’ behavior during navigation. For example, a speed limit sign can cause a participant to reduce their speed. Animacy may therefore reflect an object’s ability to influence participants’ behavior rather than strictly biological agency.

One minor confound in this study is that statistics of the “ground” label differed between the movie-watching experiment and the navigation experiment. In the movie watching experiment, the movies were hand-tagged by an observer who selected two to four of the most salient object categories that appeared in each second of the movies. Inspection of the tags revealed that the ground was not always labelled, despite being present in many scenes. Instead, the ground was often labeled with environmental descriptors such as “tundra” or “savannah,” that co-occurred with animals. This bias likely caused the animacy analysis to show that the ground was represented as an animate category in the movie-watching experiment (Fig. 5A). This bias did not arise in the navigation task, as semantic segmentation was performed by the game engine, which separated ground labels from animate agents. The animacy of the ground category should therefore be interpreted with caution in the movie-watching experiment due to this labeling confound.

The data for the two tasks were collected from separate groups of participants. To calculate the tuning shifts between tasks, the group-average weights from the movie-watching participants were mapped to the navigation participants’ cortical surfaces for comparison. Previous studies from this lab had shown that first 5-6 semantic PCs are consistent across participants (Huth et al., 2012, 2016; Deniz et al., 2019), and that the group-average model explains about 78% of the variance in individual participants (Visconti di Oleggio Castello et al., 2025). These results suggest that the movie-watching group model provides a reasonable baseline for navigation participants. However, some individual variance is inevitably lost through group averaging and cross-subject mapping, potentially reducing the sensitivity of these analyses to tuning shifts within individual participants. Future studies using the same participants across both tasks will provide greater sensitivity for characterizing task-related tuning shifts within individuals.

The results reported here extend previous work that examined how attention shifts semantic tuning across the human cortex. While previous studies have shown that attention shifts semantic tuning in the human brain towards attended semantic concepts (Çukur et al., 2013; Harel et al., 2014; McKee et al., 2014; Nastase et al., 2017; Shahdloo et al., 2022), tunings shifts between different tasks has not been well-characterized. Here we show that semantic tuning in the human brain shifts between movie-watching and active navigation, two naturalistic tasks with distinct behavioral demands. These results demonstrate that the human brain dynamically optimizes its limited neural resources towards task-relevant information, enabling it to efficiently perform diverse tasks across complex, ever-changing environments.

## Acknowledgments

We thank all members of the Gallant Lab for discussion and support throughout this work, especially A. Nunez-Elizalde, M. Lescroart, J. Gao, M. Eickenberg, S. Slivkoff, and S. Popham. We also thank Dimitar Filev and Ken Washington at the Ford Motor Company, and Marc Steinberg at the Office of Naval Research, who provided critical funding support during the early stages of this project.

## Funding

National Institutes of Health grant R01-EY031455 (JLG)

Office of Naval Research grant N000142012002 (JLG)

Ford University Research Program (JLG)

Office of Naval Research DURIP awards N00014-22-1-2217 (JLG)

Office of Naval Research Multidisciplinary University Research Initiative grant N000141410671 (JLG)

National Science Foundation Graduate Research Fellowship Program DGE 1106400 & DGE 1752814 (TZ)

## Author contributions

Conceptualization: TZ, JLG

Methodology: TZ, JLG

Investigation: TZ

Formal Analysis: TZ,

Writing – original draft: TZ

Writing – review & editing: TZ, JLG

## Conflict of interest

The authors declare no competing financial interests.

## Notes

### Competing Interest Statement

The authors have declared no competing interest.

### Summary of Updates

This fixes the broken PDF conversion from version 1 that resulted in low-resolution figures and an un-rendered equation.

