## Supplemental figures 1-10 for "Visual semantic tuning across the cortex shifts between tasks"

The file includes:

Supplemental figures 1-10

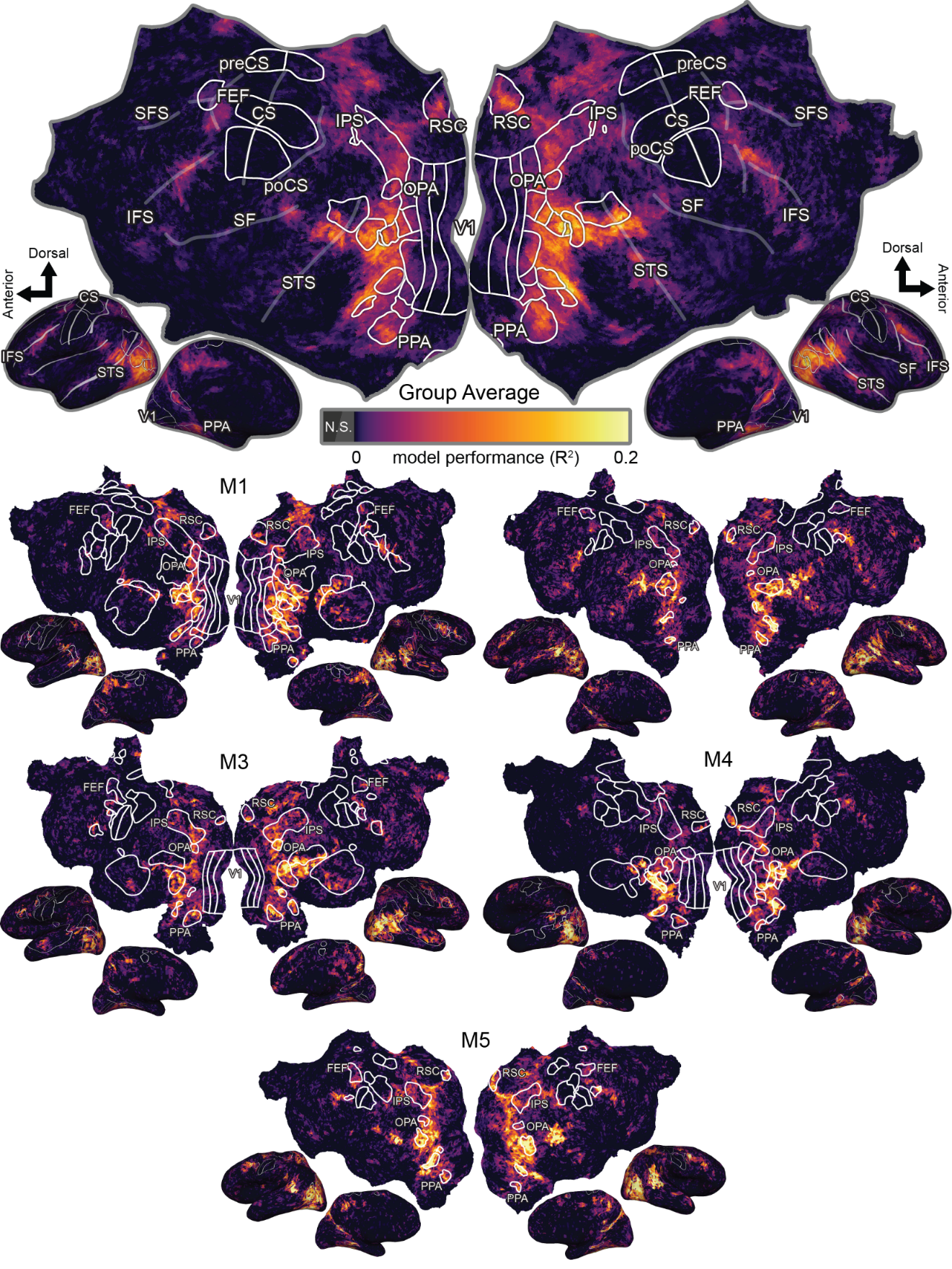

Supplemental Figure 1. Visual semantic model performance in movie-watching. Voxel are colored by the split R^2^ model performance for the visual semantic model. The group-average performance is shown at the top on the fsaverage surface, and individual subjects are shown below the group average. Visual semantics explains activity across the anterior visual cortex, including RSC, IPS, OPA, MT, LO, FFA, and PPA, and also parietal regions such as the precuneus and TPJ, and also parts of the precentral sulcus.

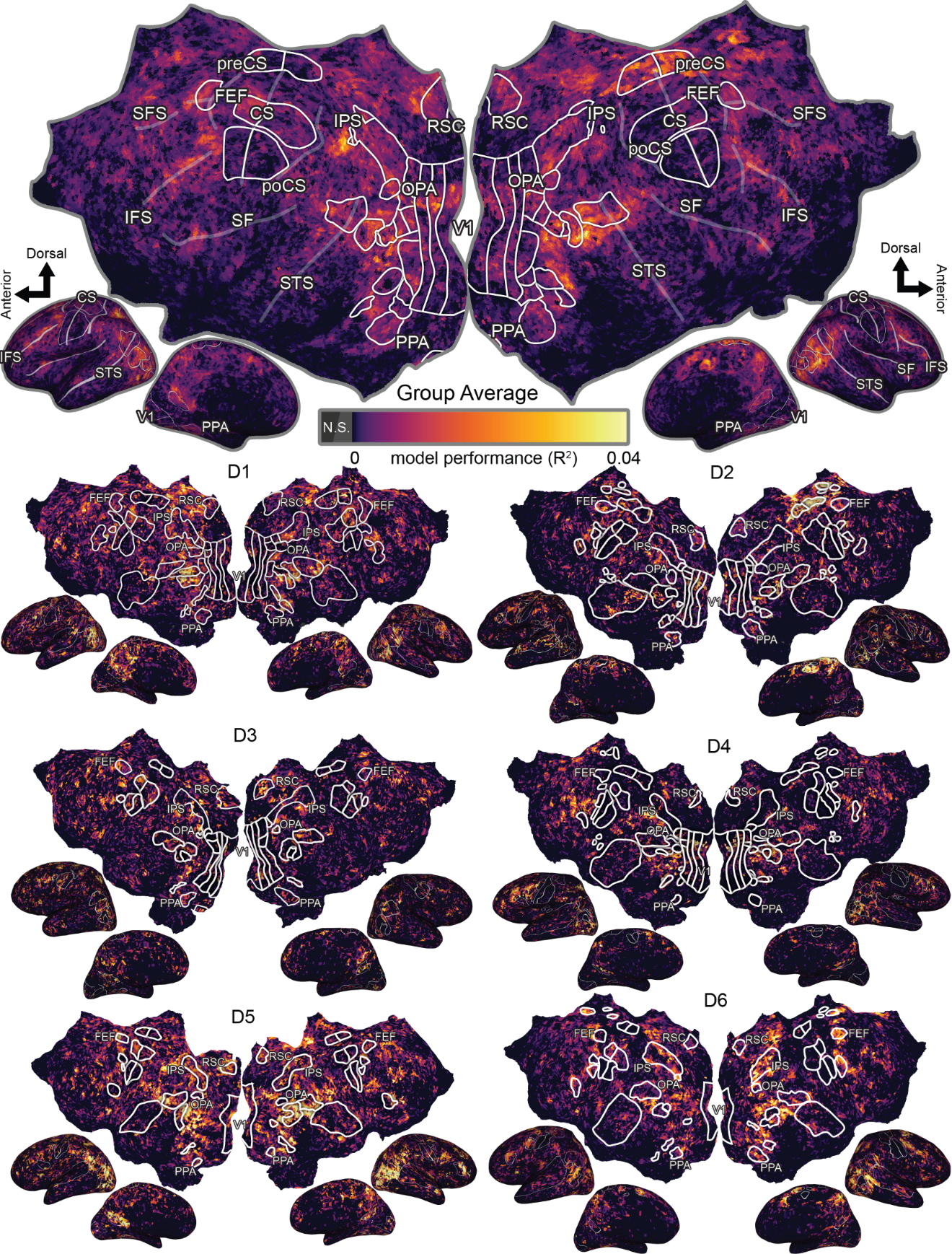

Supplemental Figure 2. Visual semantic model performance in driving. Voxel are colored by the split R^2^ model performance for the visual semantic model. The group-average performance is shown at the top on the fsaverage surface, and individual subjects are shown below the group average. Visual semantics explains activity across the anterior visual cortex, including RSC, IPS, OPA, and PPA, and also parietal regions such as the TPJ. Additionally, the visual semantic model also explains activity in multiple prefrontal regions outside known ROIs, including the posterior medial PFC and along the SFS and IFS, suggesting that the representation of visual semantic information may be more distributed across the cortical surface during driving than movie-watching.

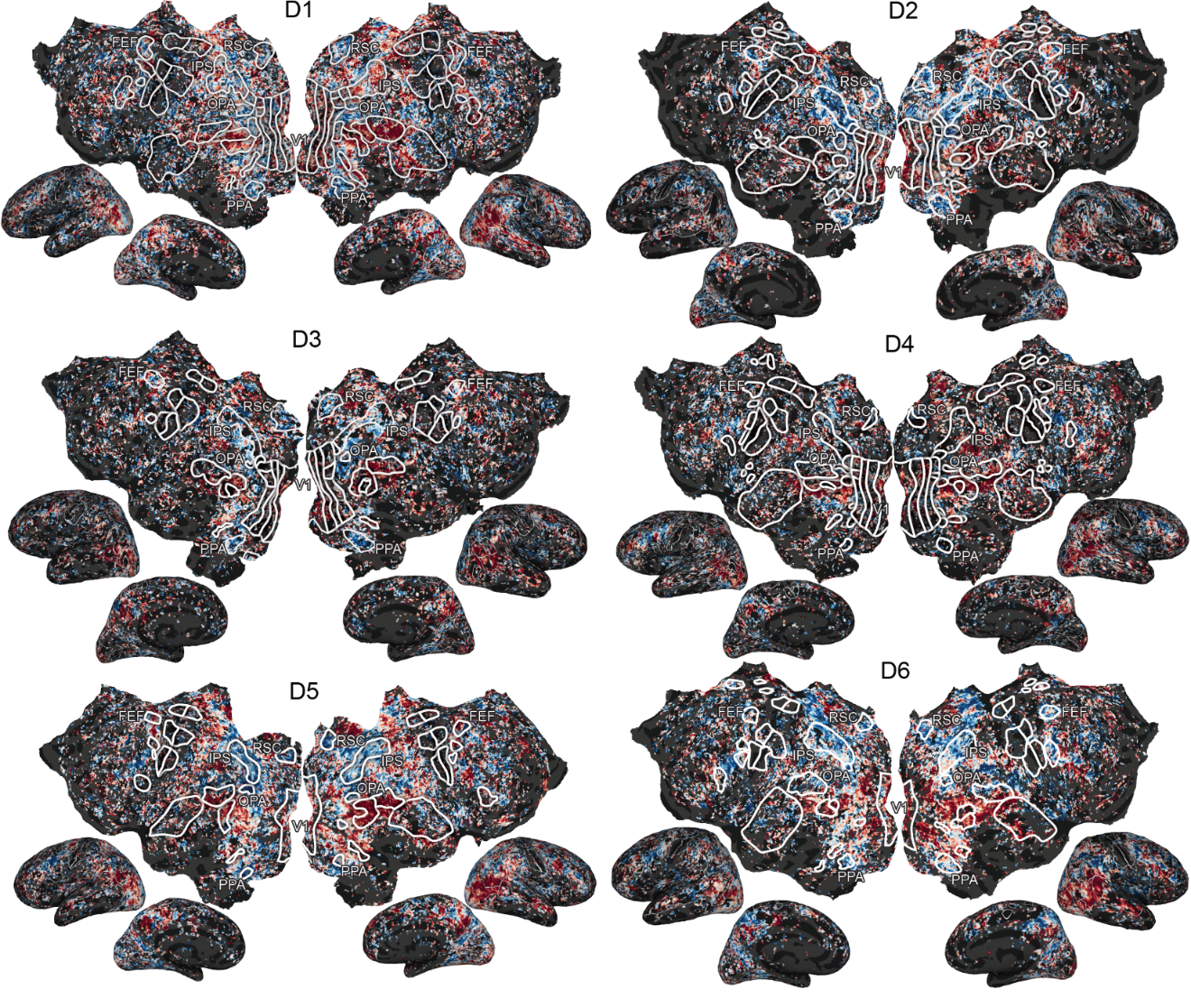

Supplemental Figure 3. Individual driving subject visual semantic tuning correlations between movie-watching and navigation as in Fig. 2A. Much of the cortex significantly shifts its visual semantic tuning between tasks. Consistent across subjects, RSC, IPS, OPA, and PPA show the strongest shift, while FFA, EBA, and EPJ show the weakest shift.

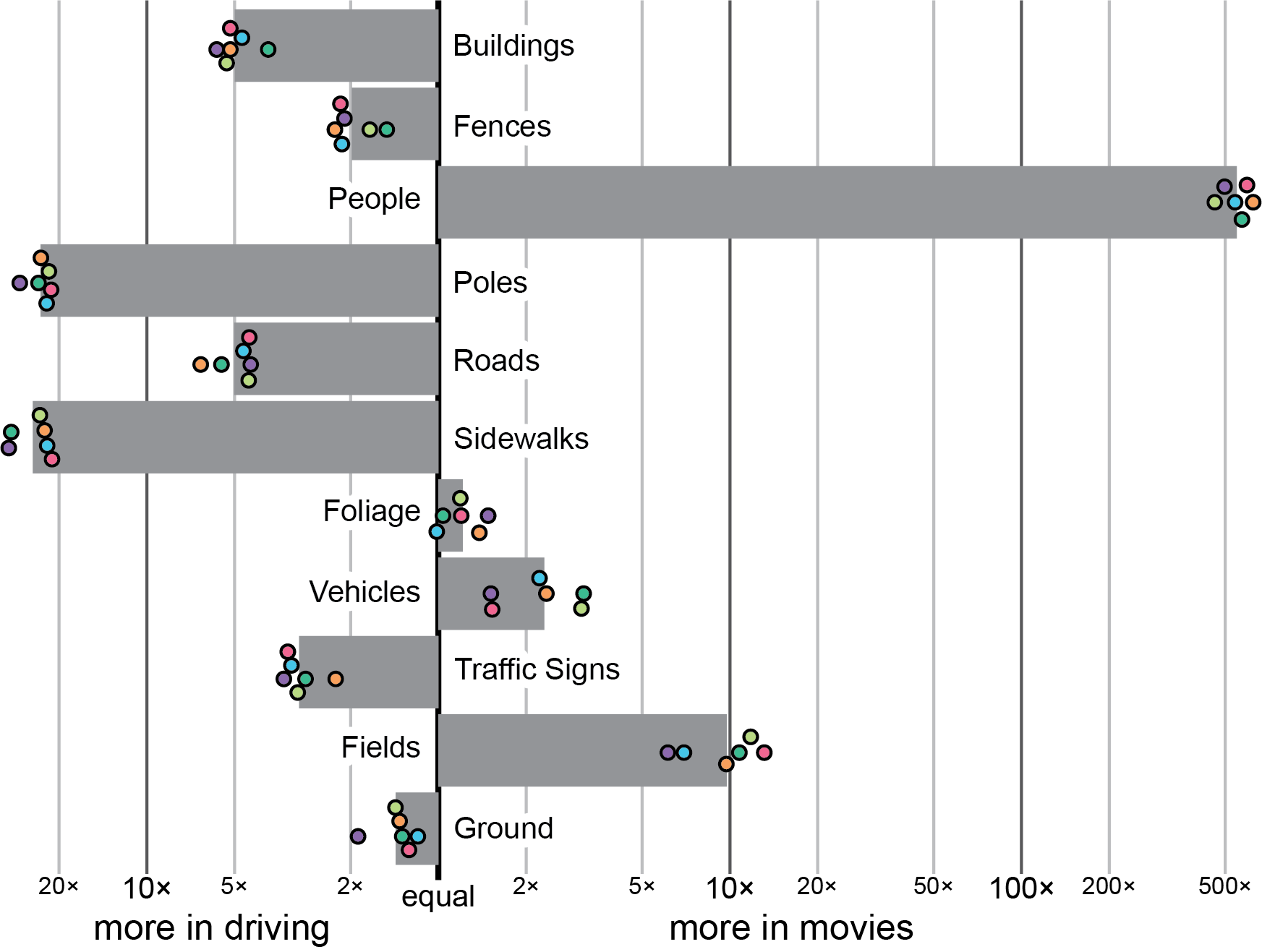

Supplemental Figure 4. Relative stimulus differences between experiments. Categories to the left of the “equal” line appeared more frequently in the navigation experiment, and those to the right of the “equal” line appeared more frequently in the movie-watching experiment. Bars show average frequency ratio over subjects, and dots indicate difference in individual subjects. These stimulus differences are not significantly correlated with any of the tuning shift components (p > 0.05, permutation test), suggesting that the tuning shifts are unlikely to be driven by the stimulus differences.

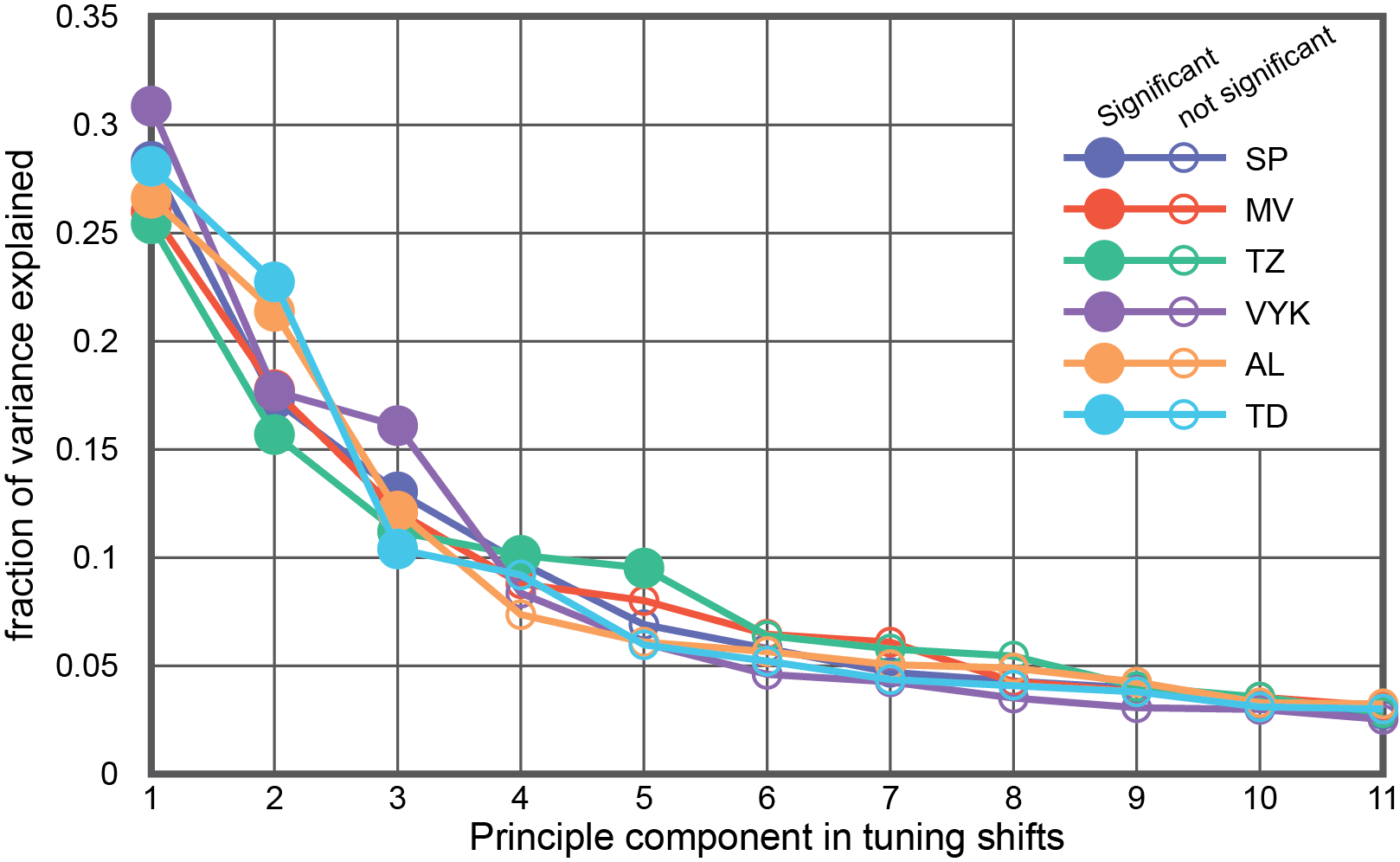

Supplemental Figure 5. Fraction variance explained by the group-level tuning shift principal components in individual subjects. Closed circles indicate components that explain a significant amount of variance (permutation test, p < 0.05), and open circles indicate those that do not. The first three PCs explain significant amounts of variances in all subjects, suggesting that there are three consistent dimensions in the cortical tuning shift between movie-watching and navigation.

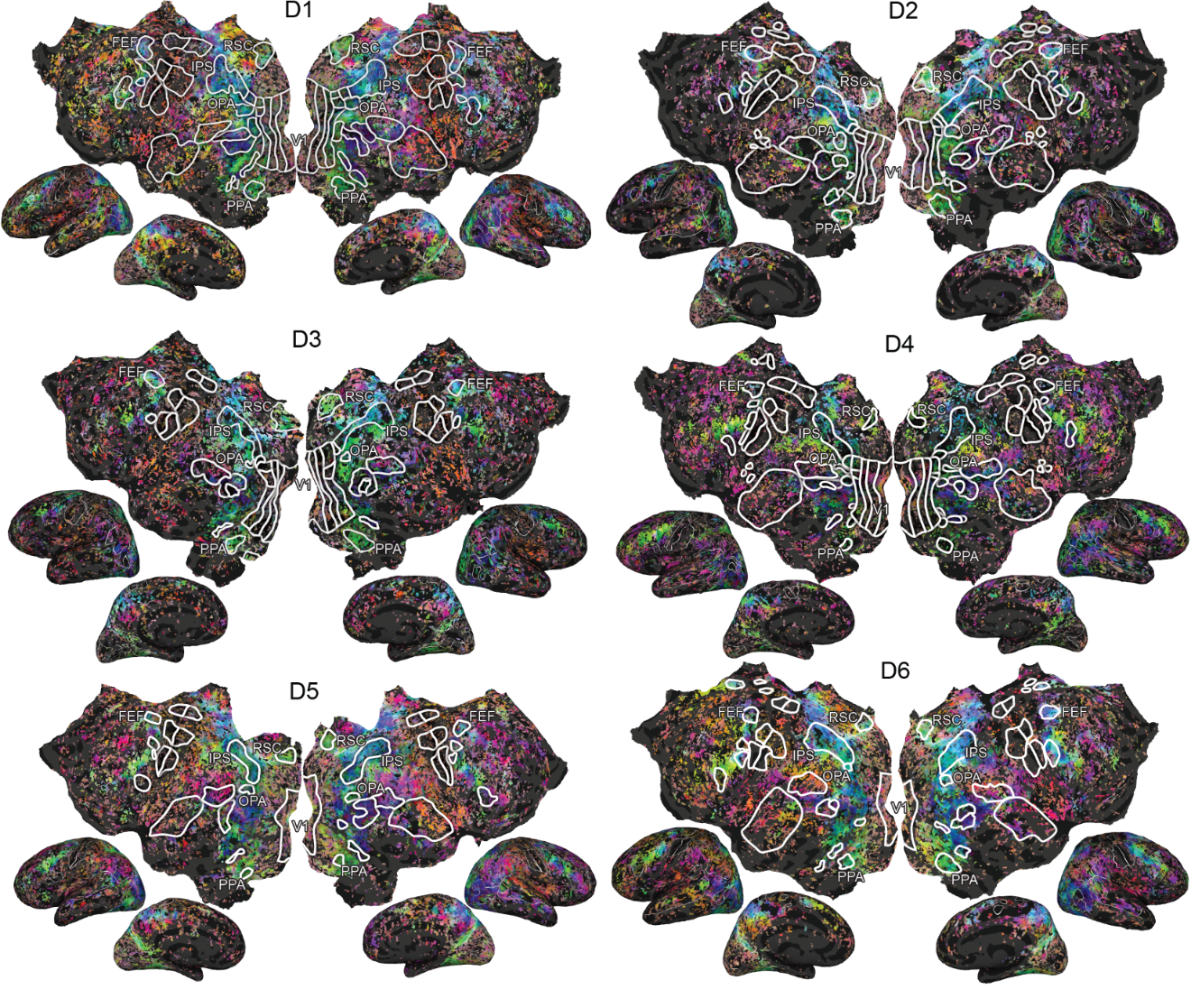

Supplemental Figure 6. Individual subject visual semantic tuning shifts in driving as shown in Fig. 3A.

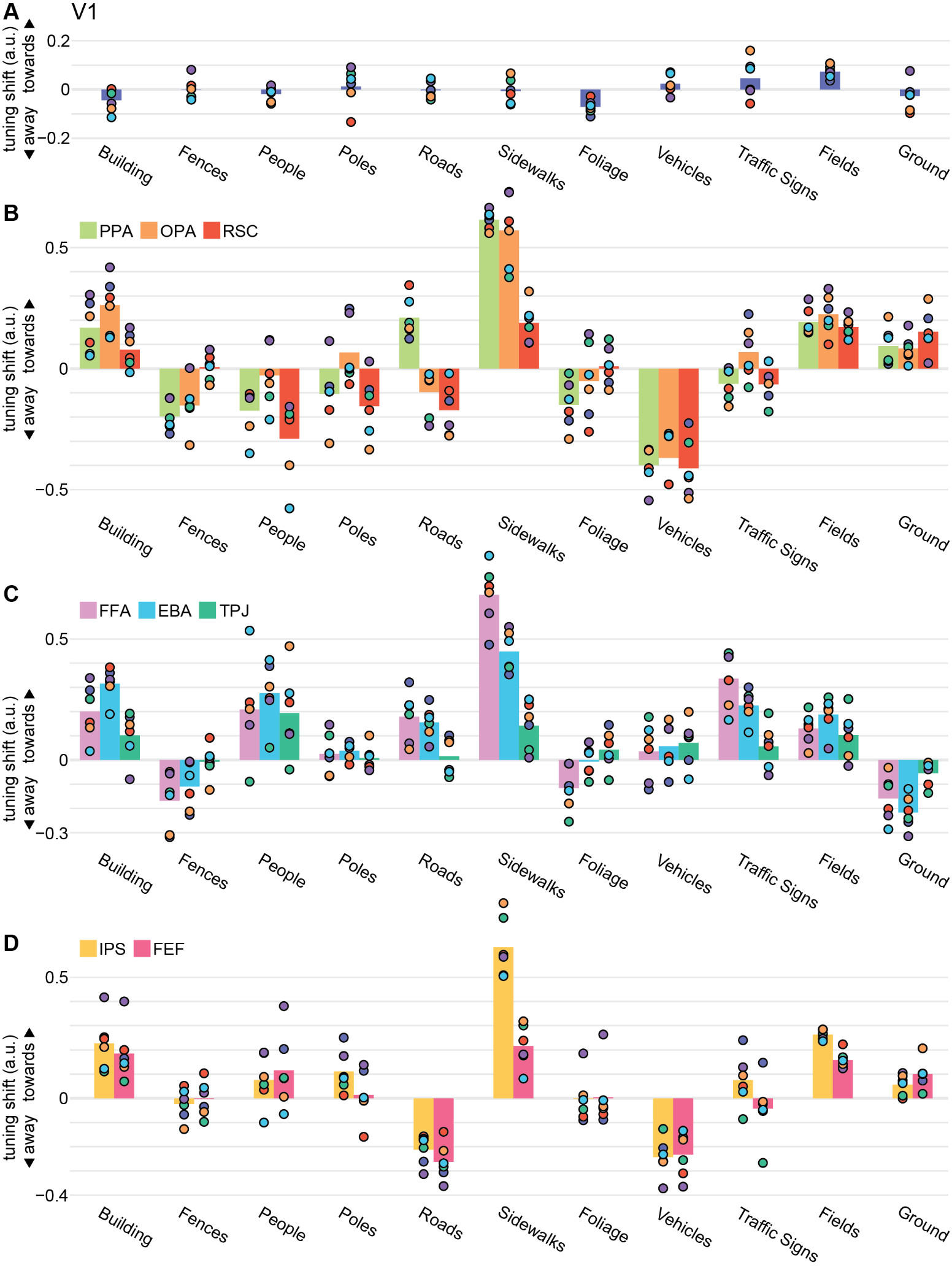

Supplemental Figure 7. Average tuning shifts in individual ROIs. The average shift (a.u.) for each category is shown for A) V1, as a comparison, B) the scene-selective regions of PPA, OPA, and RSC, C) people- and socially-selective regions of FFA, EBA, and TPJ, and D) visual attention regions of IPS and FEF. Bars show average shift across subjects, and dots indicate individual subjects.

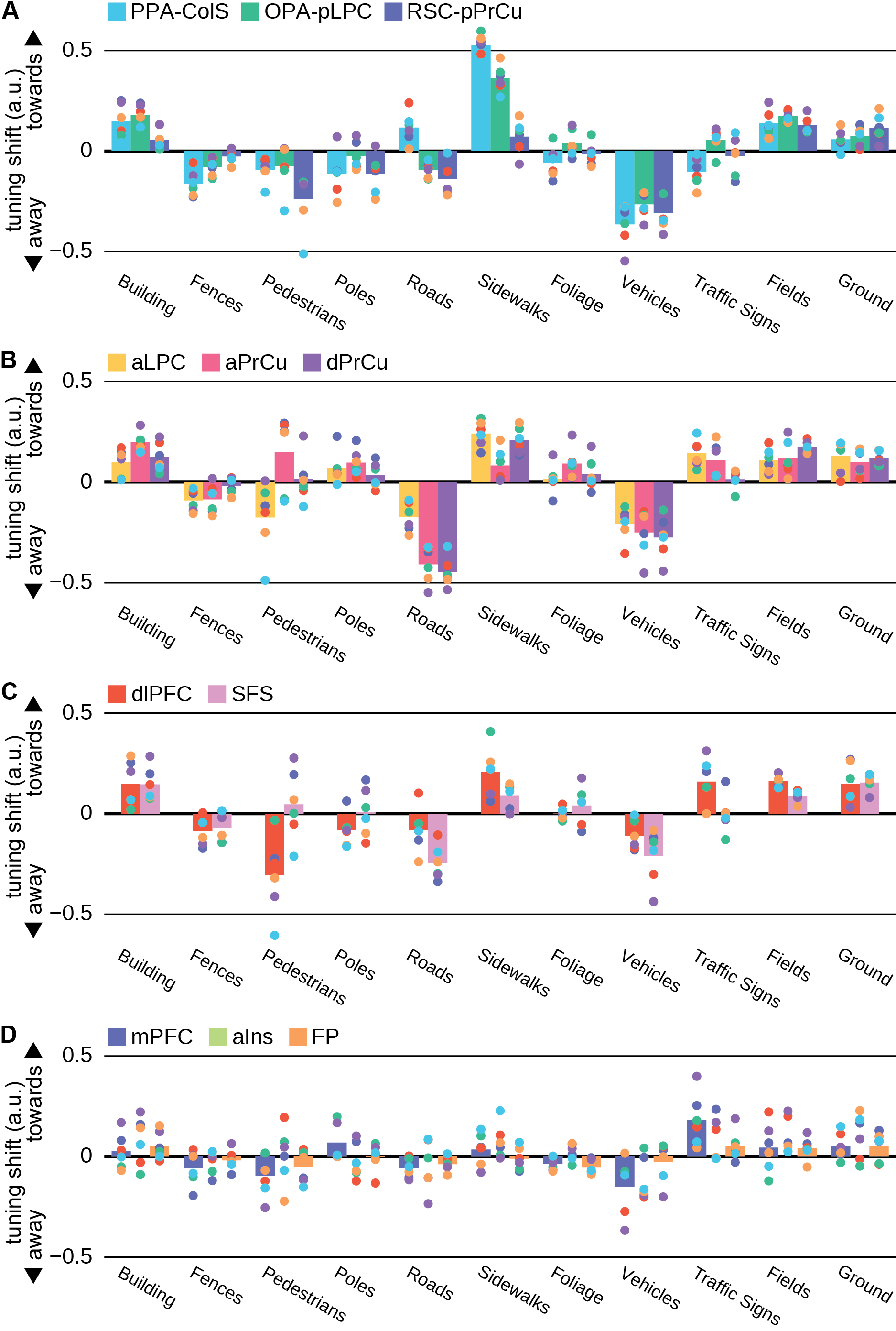

Supplemental Figure 8. Average tuning shifts in the 11 individual ROIs in the cortical navigation network. The average shift (a.u.) for each category is shown for A) PPA-ColS, OPA-pLPC, and RSC-pPrCu, B) aLPA, aPrCu, and dPrCu, C) dlPFC and SFS, and D) mPFC, aIns, and FP. Bars show average shift across subjects, and dots indicate individual subjects.

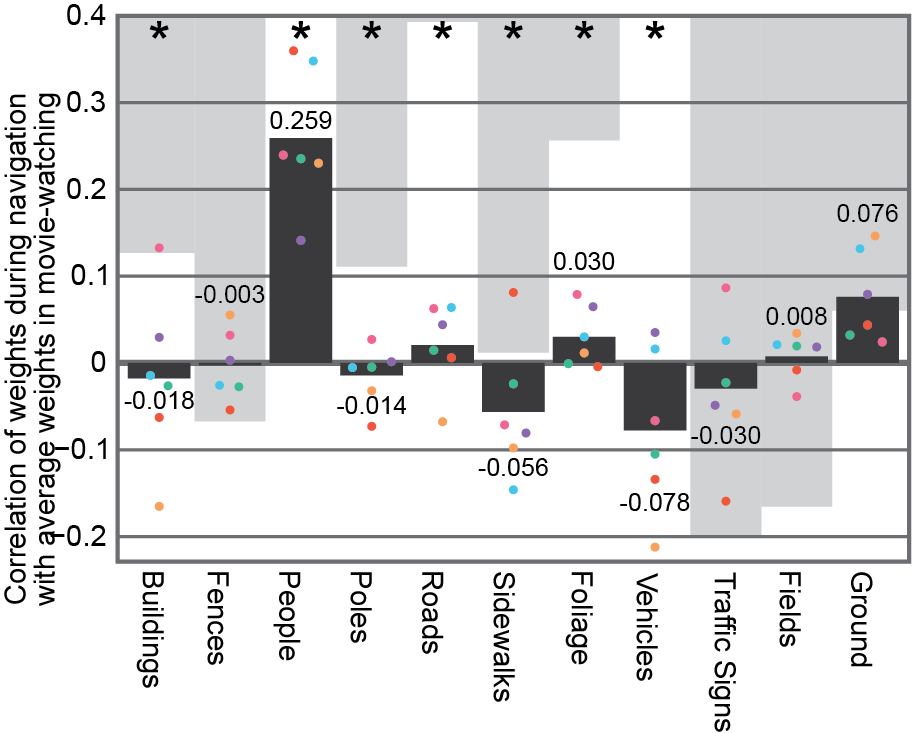

Supplemental Figure 9. Object representations in the cortex differ across tasks. As the cortex shifts its tuning, object categories become represented by different cortical networks. To quantify this difference, we computed the correlation between the weight vectors for each of the object categories between the two tasks. Bars show the group-average correlations for each category, and dots indicate individual subjects. Shaded regions indicate 95% confidence interval for the null distribution that there is no difference in the representation of each object category between navigation and movie-watching (for pedestrians and vehicles, the bottom of this confidence interval is above the upper bound of the plot). Stars indicate object categories that are significantly difference between the two tasks at the group level (FDR corrected p < 0.05, bootstrap test). The representations of buildings, pedestrians, poles, roads, sidewalks, foliage, and vehicles were significantly different between the two tasks. The difference in representation is not constant across categories: the representation of pedestrians is most similar between tasks, and the representation of vehicles is most dissimilar. These results suggest that task-related tuning shifts induce differences in the representations of object categories, and that these differences may be category-specific.

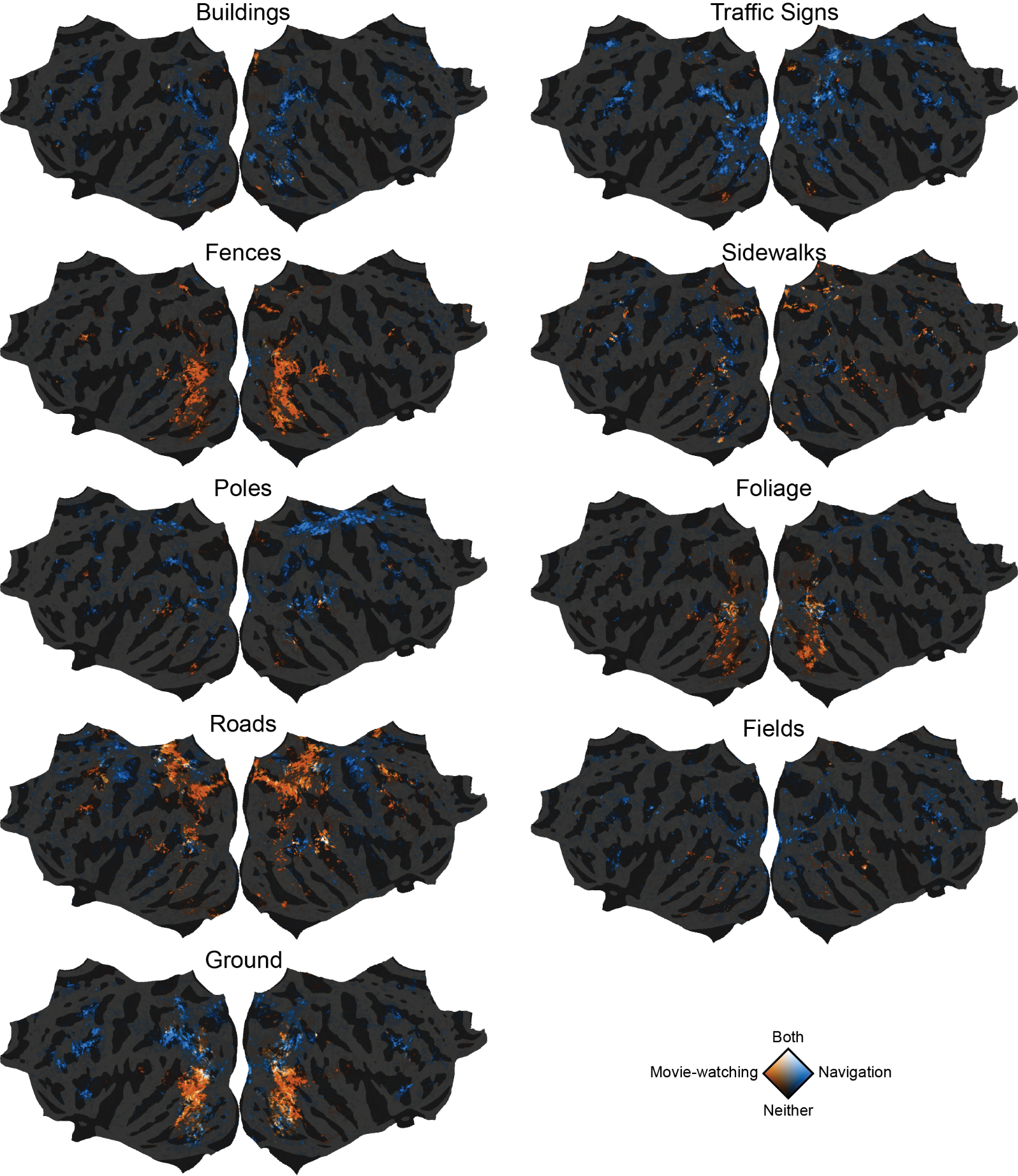
Supplemental Figure 10. Weight comparison for other object categories. Color scheme is the same as in Fig. 5C and D
